# Model-based deconvolution for DSC-MRI: A comparison of accuracy, precision, and computational complexity of parametric transit time distributions

**DOI:** 10.1101/2023.02.12.528216

**Authors:** Rashed Sobhan, Pinelopi Gkogkou, Glyn Johnson, Donnie Cameron

## Abstract

**Object:** Dynamic susceptibility contrast MRI (DSC-MRI) is the current standard for cerebral perfusion estimation. Model-dependent approaches for DSC-MRI analysis involve assuming a parametric transit time distribution (TTD) to characterize the passage of contrast agent through tissue microvasculature. Here we compare the utility of four TTD models: namely, skewed-Gaussian, gamma, gamma-variate, and Weibull, to identify the optimal TTD for quantifying brain perfusion.

**Materials and Methods:** DSC-MRI data were acquired in nine subjects at 1.5T, and normal-appearing white- and gray-matter signals were assessed. TTDs were compared in terms of: goodness-of-fit, evaluated using RMSE; noise sensitivity, assessed via Monte-Carlo-simulated noisy conditions; and fit stability, quantified as the proportion of total fits converging to the global minimum. Computation times for model-fitting were also calculated.

**Results:** The gamma TTD showed higher fit stability, shorter computation times (*p*<0.008), and higher robustness against experimental noise as compared to other models. All functions showed similar RMSEs and the parameter estimates (*p*>0.008) were congruent with literature values.

**Discussion:** The gamma distribution represents the most suitable TTD for perfusion analysis. Moreover, due to its robustness against noise, the gamma TTD is expected to yield more reproducible estimates than the other models for establishing a standard, multi-center analysis pipeline.

## 1. Introduction

Dynamic susceptibility-contrast magnetic resonance imaging (DSC-MRI) is currently the standard method for the quantification of cerebral perfusion. It provides clinicians with crucial information for diagnosis, assessment, treatment planning, and monitoring of several cerebral pathologies, such as tumors [1], ischemic stroke [2], multiple sclerosis [3], and Alzheimer’s disease [4]. The imaging technique involves acquiring a series of rapid T2 or T2*-weighted images during the first pass of a Gadolinium-based contrast agent (GBCA) bolus [5]. Signal changes are converted to estimates of contrast concentration, from which cerebral hemodynamic parameters such as cerebral blood flow (CBF), cerebral blood volume (CBV), and mean transit time (MTT) can be calculated.

The fundamental equation used in DSC-MRI is: [6,7]

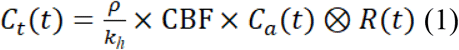

where *C_t_*(*t*) is tissue GBCA concentration as a function of time; *ρ* is the brain density; *k_h_* is a dimensionless constant accounting for the difference in hematocrit between capillaries and large arteries; *C_a_*(*t*) is the GBCA concentration in the artery feeding the tissue (also known as the arterial input function, AIF); *R*(*t*) is the tissue residue function, which represents the fraction of tracer entering the tissue at time zero that remains at time *t*; and ⊗ represents convolution. Values of *ρ* and *k_h_* have previously been measured as 1.04 g/ml and 0.73, respectively [6–8].

The product of CBF and *R*(*t*) is obtained by deconvolving *C_t_* with *C_a_*. Since, by definition, *R*(0) is equal to one, CBF can be determined as the initial value of the product CBF×*R*(*t*)[9,10]. The residue function can be used to derive the tissue transit time distribution (TTD), *h*(*t*): namely, the density function representing the range of transit times whereby contrast particles traverse capillary tubes of different lengths in a given region of interest (ROI) [9,11]:

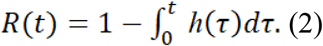

The MTT is given by:

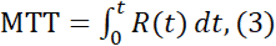

and CBV can then be calculated using the central volume principle [12]:

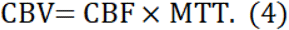

Deconvolution is an ill-posed problem; small noise variations in *C_a_* can cause large oscillations in the resulting estimate of *R*. Deconvolution must thus be constrained in some way to produce physically meaningful estimates of CBF and *R*. Two general approaches have been proposed: model independent and model dependent.

The model independent approach involves solving equation (1) using a transformation approach based on the convolution theorem [8,13,14] or an algebraic approach, where equation (1) is expressed as a series of simultaneous equations and solved by singular value decomposition (SVD) [10,15–17]. In either case the process must be subject to some form of filtering or regularization to provide a stable solution. The SVD approach is now the standard, although it is subject to several obvious problems. First, even with quite sophisticated forms of regularization (e.g., Tikhonov [10]) the resulting residue function is often unrealistic, with negative values and submaximal *R* at *t* = 0. Moreover, the resulting errors in flow estimates are sensitive to the degree of regularization employed [18,19]. These errors are exacerbated if there are time delays between *C_a_* and *C_t_*, though these errors can be reduced by using block circulant SVD deconvolution [17].

The model dependent approach assumes a parametric model for *R*. The solution of equation (1) then reduces to a least squares minimization problem. Several studies [6,20–22] have adopted this approach, assuming physiologically plausible forms for *h*(*t*) and fitting trial values of *R*, as calculated by equation (2), to equation (1). Physiologically plausible functions for *h* and *R* must conform to several constraints. Blood cannot traverse the tissue instantaneously, and contrast must eventually exit the tissue. So, *h*(0) = 0 and *R* (0) = 1; as *t* ⇒ ∞, *h* ⇒ 0, *R* ⇒ 0; and its integral should be unity. In addition, *R* should be smooth and monotonic, and it should decrease as a function of time[6].

Although several different functions for *h* (and hence *R*) have been proposed [6,20,22], no study, to our knowledge, has compared these functions in terms of their computational benefits and robustness against noise. In this study, we compare the relative utility of three already-published forms of *h*(*t*), and a newly proposed form, with a view to identifying the most suitable model for future clinical use.

## 2. Theory

### Skewed-Gaussian TTD

To represent the underlying asymmetry in *h*(*t*), Koh *et al.* proposed a skewed-Gaussian TTD with the following form:

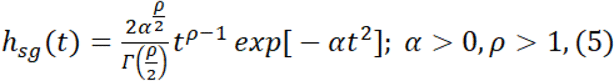

where *α* is the scale parameter and *ρ* is the shape parameter [20]. The shape parameter *ρ* is bounded at unity to avoid an exponential *R*. The gamma function of equation (5) is defined as follows:

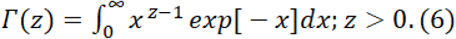

### Gamma TTD

Mouridsen et al. suggested that a family of gamma distributions with certain constraints can plausibly describe blood flow through healthy [6,21] and ischemic [23] tissue vasculature. It has also been used in studies on the interdependence of the cerebral metabolic rate of oxygen and CBF [24,25]. The gamma TTD is given by:

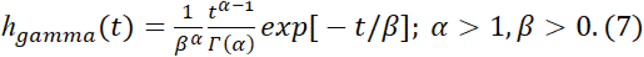

Here, *β* is the scale parameter and *α* is the shape parameter. The gamma function Γ(α) is given by equation (6). An exponential *R*(*t*) is avoided by setting *α* > 1, and an infinite value of gamma *h* is avoided by setting *β* > 0.

### Gamma-variate TTD

In a recent study, a gamma-variate distribution was used as a TTD to model dynamic contrast enhanced (DCE)-MRI data [22]. The usual form for the gamma variate function is given by:

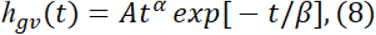

where *A, α*, and *β* are free parameters [26].

Madsen proposed a normalized gamma-variate function by decoupling the parameters, thereby making it more robust for least-squares fitting [26]. The form suggested by Madsen is as follows:

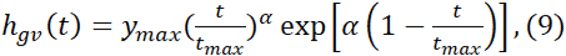

where *y_max_* is the maximum value of the distribution, *α* is the decay parameter, and *t_max_* is the time at which *h_gv_* is maximum. The unit integral constraint of *h* yields:

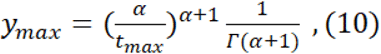

where Γ is the gamma function defined by equation (6) [27,28]. Substituting *y_mca_* from equation (10) into equation (9) and substituting *t’* gives:

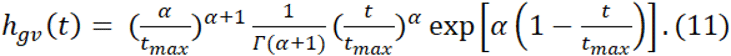

### Weibull TTD

The Weibull distribution is widely used for modeling reliability data, hazard functions, failure times, and analyzing the lifetimes of electrical as well as mechanical components [29]. Previously, it has been used to represent the distribution of capillary transit times for modelling the oxygen extraction fraction [24]. The normalized form of the Weibull distribution is given by:

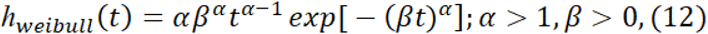

where *α* is the shape parameter and *β* is the reciprocal of the scale parameter, *η* (i.e. *β* = 1/*η*).

Our rationale behind exploring the Weibull distribution as a candidate TTD function is as follows. First, it conforms to the constraints for a TTD to be representative of physiologically-realistic vasculature. Second, it can transform into several different distributions by varying *α*. When 0 < *α* < 1, the TTD decreases exponentially from an infinite initial value. These values of *α* are avoided here as the constraint *h*(0) = 0 is violated; a value of *α* = 1 produces an undesired exponential *R*(*t*). With 1 < *α* < 2, the TTD rises sharply with slow rate of fall (positively skewed). With *α* = 2, it becomes a Rayleigh distribution, and in the range 3 < *α* < 4 it becomes a symmetrical, bell-shaped curve, resembling a Gaussian distribution, starting at *t* = 0. At higher values of *α* (> 10), it takes the shape of an extreme value distribution, which is negatively skewed [30]. Therefore, we speculate that the application of Weibull TTD can facilitate characterization of a variety of microvascular environments, observed in both normal and pathologic tissue.

All four TTDs used above are right-skewed, resembling the underlying asymmetry in the physical system and the distribution of transit times [20]. Figure 1 gives examples of these four TTDs in three different tissue conditions: healthy gray matter (GM), ischemic, and tumor. The parameter values for simulating these TTDs were taken from published studies [21,22,31,32] (see Supporting Information).

**Fig 1.**
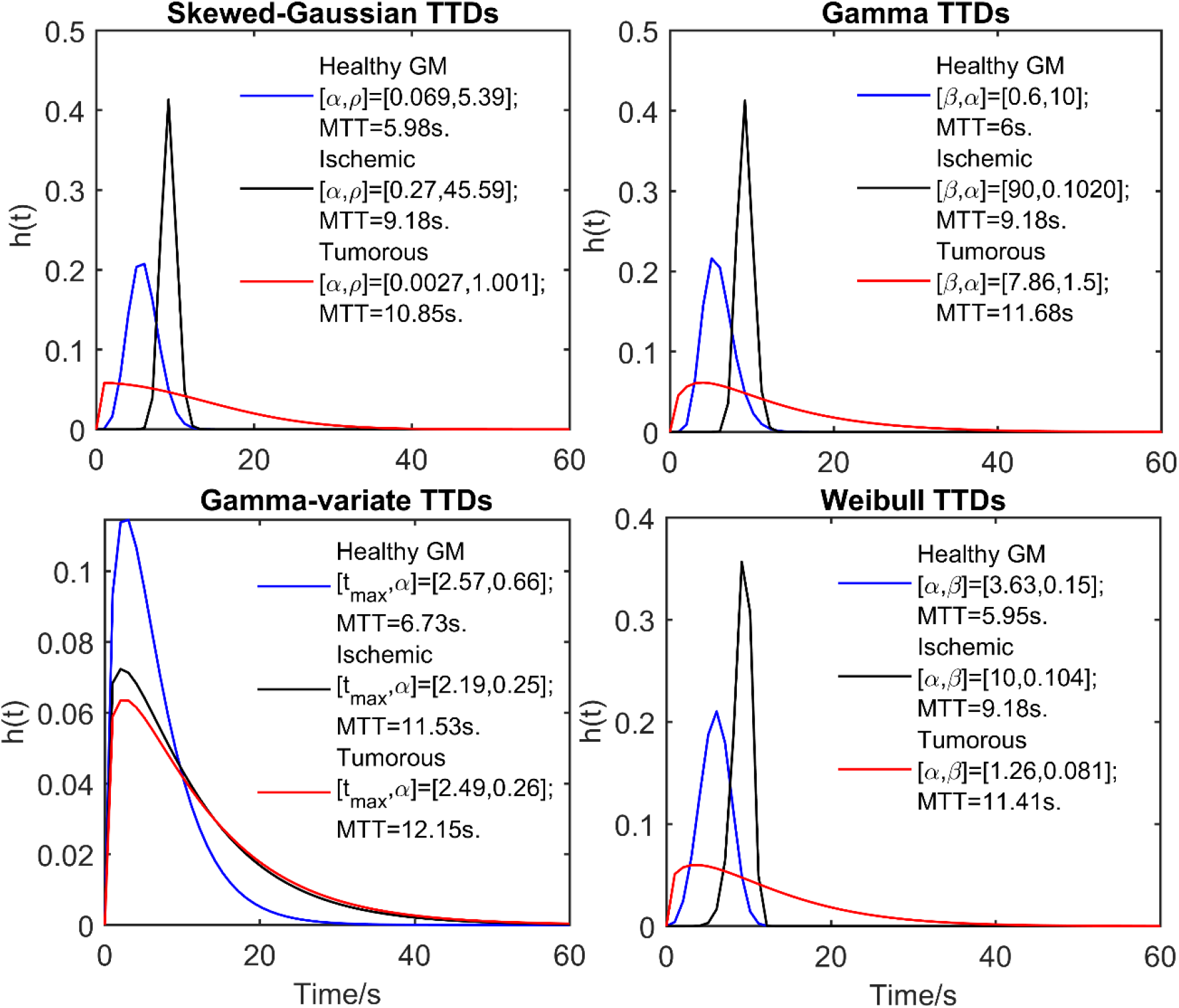
(Clockwise, starting from top left) Simulated skewed-Gaussian, gamma, Weibull, and gamma-variate TTDs in healthy (gray matter), ischemic (internal carotid stenosis) [22], and tumor (high grade glioblastoma multiforme) [21] conditions, respectively in blue, black and red. Parameter values for the gamma TTD were taken from the Dynamic Contrast-enhanced (DCE)-MRI studies of Schabel [21] and Larsson et al.[22], whereas parameters for all other TTDs were measured through curve-fitting of simulated signals (see Supporting Information for details)

## 3. Methods

### 3.1. Data acquisition

The data were derived from DSC-MRI scans of nine low-grade glioma patients from a previously-published, Institutional Review Board approved study [33]. The DSC-MRI data were acquired at 1.5 tesla (Siemens Vision/Symphony; Siemens Healthineers, Erlangen, Germany) with a gradient-recalled-echo echo-planar imaging sequence during the first pass of a standard dose (0.1 mmol/kg) bolus of gadopentetate dimeglumine (Magnevist, Berlex Laboratories, Wayne, NY). Imaging parameters were as follows: Repetition time (TR) = 1000ms; Echo time (TE) = 54 ms; field of view = 230 × 230 mm; section thickness = 5 mm; matrix size = 128 × 128; in-plane voxel size = 1.8 × 1.8 mm; interslice gap = 0–30%; flip angle = 30°; and signal bandwidth = 1,470 Hz/pixel; total no of slices = 7 or 10. Contrast was injected at a rate of 5 ml/sec, followed by a 20 ml bolus of saline at 5 ml/sec. A total of 60 images were acquired at one second intervals, giving a total acquisition time of one minute. The contrast injection coincided with the acquisition of the fifth image in the series, meaning the bolus would typically arrive at the fifteenth to twentieth image.

### 3.2. Image analysis

All analyses were performed offline using a Windows workstation (2.50–2.70 GHz dual-core Intel^®^ core™ i5-7200U CPU, 8GB RAM).

#### Region of interest selection

For each subject, signal time curves (STCs) were obtained from 3×3-pixel ROIs placed manually in four areas of normal-appearing white matter (WM) in the frontal and parietal lobes and in two areas of GM in the caudate nucleus, as marked in Figure 2 (inset). Representative average signals from the marked GM and WM ROIs are shown in Figure 2. The short acquisition duration and ROIs planned in healthy brain tissue (see below) allowed us to ignore GBCA extravasation through the blood brain barrier while modelling the TTDs. All processing and analysis of the acquired signals was performed offline with code written in MATLAB (R2018a, Natick, MA).

**Fig 2.**
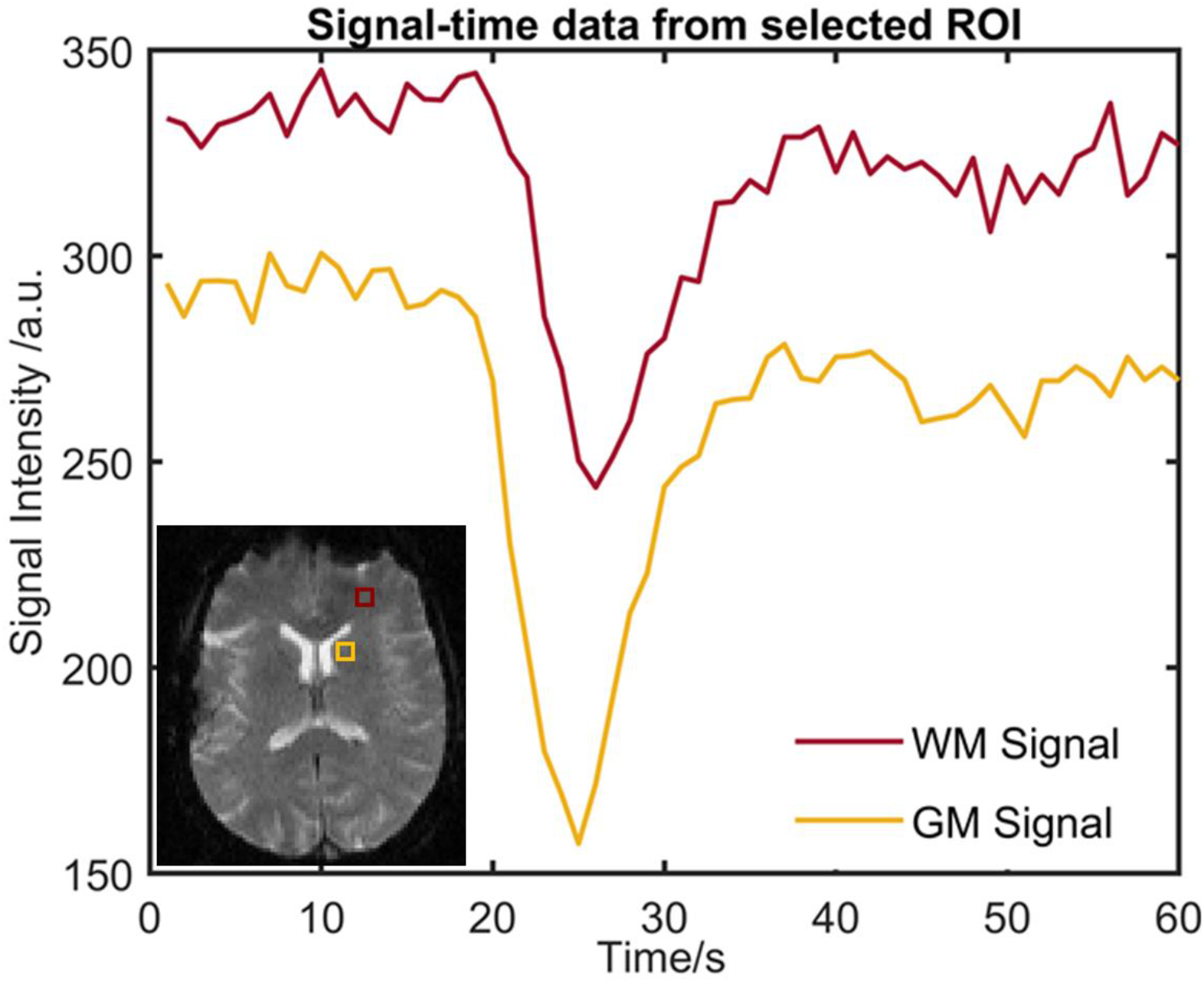
Average DSC-MRI signal intensity curves (in arbitrary units, a.u.) for the white matter (WM) and gray matter (GM) regions of interests (ROIs) (in maroon and yellow, respectively) placed on the first time point of a gradient-recalled-echo (GRE) image. Inset: The pairs of 3 × 3-ROI were placed in the caudate nucleus for GM (shown with yellow square) and the frontal lobe for normal appearing WM (shown with maroon square)

#### Arterial input function detection

Arterial voxels (AVs) were automatically identified following the processes used in several published studies [7,34,35]. After background voxel removal and skull stripping, each brain STC was converted into a concentration time curve (CTC) using the equation below [27,28]:

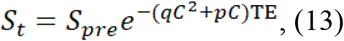

where *q* and *p* are constants that depend on the external magnetic field strength, *B_0_*, and TE is the echo time of the acquisition pulse sequence. At a *B_0_* of 1.5 T, *q* = 0.74 s^-1^mM^-2^ and *p* = 7.2 s^-1^mM^-1^. As per routine MRI perfusion processing, the first six images were discarded, as they did not demonstrate an equilibrium signal for the baseline calculation [36]; this was also confirmed by visual inspection of STCs. *S_pre_* is therefore calculated as the mean of the signal intensity values from the seventh image till the image preceding the bolus arrival.

Motion- and partial-volume-affected CTCs were discarded using a threshold on roughness index, given as integral of the second derivative of the CTC with respect to time. Twenty five percent of the CTCs with the highest RI were discarded, as per previous studies on automatic AIF detection [7,34]. We then leveraged the fact that soft-tissue CTCs are typically wider than AIFs, with lower peak concentration and area under the curve (AUC), and slower washout, to remove additional spurious CTCs. To this end, multiple thresholds were applied to different CTC features: 50% of the CTCs with the highest full width at half maximum (FWHM) and 20% with the highest first moment (FM) were removed. Then we rejected 30% of the remaining CTCs with the lowest peak concentration, and lastly, 40% of the CTCs with the lowest area under the curve (AUC) [8,34,35].

These less-stringent thresholds identified all the true AVs, but at the cost of also identifying many false AVs. To group the true AVs, the thresholding step above was followed by standard *k*-means clustering[37]. This divided the remaining CTCs into five clusters, putatively corresponding to GM, WM, arterial blood, venous blood, and ‘other’, such as ventricles containing cerebrospinal fluid (CSF) [7,34,38]. Each cluster was represented by a centroid. The final AIF was obtained by aligning and averaging the CTCs within the cluster whose centroid had the highest peak and lowest FM.

#### Curve fitting and parameter estimation

For each tissue, the parametric equation for *C_t_* was derived according to equation (1) using three free parameters: two corresponding to the shape and scale of *h* and the third to the CBF. This parametric *C_t_* was then converted to signal using the non-linear equations of Patil et al. [27] and fitted to measured tissue signal values. We chose not to convert raw STCs into CTCs, unlike in standard DSC-MRI analysis, as noise in the signal (but not concentration) is independent of amplitude. Fits were repeated for one hundred random combinations of initial guesses uniformly distributed over their likely physiological limits.

For curve fitting, MATLAB’s ‘lsqcurvefit’ least-squares algorithm was used. Optimization settings were as follows: algorithm = trust-region-reflective[39]; step and function tolerance = 1×10^-20^; maximum number of function evaluations = 5,000; maximum number of iterations = 2,000. All other settings were left at their default values. After convergence, the CBF and the TTD model parameters were obtained. MTT was calculated from (equation 3) and CBV was derived via the central volume principle (equation 4).

### 3.3. Model Comparisons

#### Goodness of fit

The goodness of fit of the estimated signals derived from each TTD model to the measured GM and WM signals was assessed via the root-mean-square error (RMSE), which was calculated as follows:

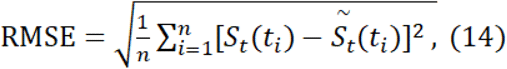

where *n* is the total number of timepoints; *S_t_*(*t_i_*) is the measured normalized signal and 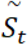, is the estimated normalized signal at the *i^th^* time point. Mean RMSE gave an estimate of the accuracy of the fits, whereas the standard deviation (SD) over multiple fits provided an indication of the precision of fits. We did not consider the model complexity along with goodness of fit, as all models had the same number of degrees of freedom.

#### Parameter Estimates

The CBV, CBF, and MTT estimates obtained with each TTD were calculated over all subjects in both GM and WM. These were compared with published literature values to assess their consistency, as were the GM-to-WM ratios for each perfusion parameter.

#### Bias and dispersion

Monte-Carlo noise simulations were used to compare the effect of noise on the accuracy and precision of perfusion estimates obtained from different TTDs. *In vivo* data were taken from a subject whose AIF was representative of most of the participant cohort. Zero-mean Gaussian noise was added in quadrature, to create a range of signal-to-noise ratios (SNRs), defined by *S_pre_/σ*, where *σ* is the SD of the pre-bolus signals. New signals were generated from the *in vivo* data for SNRs of 10-100, in steps of 10, each with 1,000 noise realizations. Parameter estimates obtained at an SNR of 100 were regarded as the “true” parameter values.

For each distribution, the relative difference between the true parameter value and the estimated parameter value averaged across different noise realizations gave a measure of bias (i.e. accuracy), whereas the standard deviation of the estimates indicated the dispersion (i.e. precision) of the perfusion parameters [40]. So,

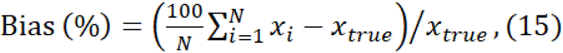

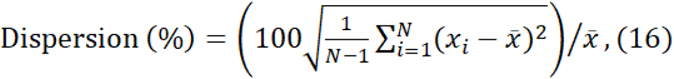

where *x_i_* is the parameter value estimated for *i*th noise realization, *x_true_* is the true parameter value, 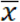 is the mean of all estimates, and *N* is the number of the noise realizations. Both bias and dispersion were calculated for all four candidate TTDs. To rank the models in terms of their accuracy and precision, the sum of the magnitude of each relative bias and dispersion estimate was taken over all SNR values, for all combinations of parameter and tissue type.

#### Success rate

To assess fit stability for each TTD, the proportion of fits converging at the global minimum was determined and is referred to here as the ‘success rate’. Models with higher success rates converged to the global minimum more frequently than other models, irrespective of the initial guess.

#### Computation time

Lastly, computation time *(T_comp_)* was calculated as the total time elapsed during the entire curve-fitting process. Means and SDs of *T_comp_* were compared for all four *h’s* over all subjects for both GM and WM.

#### Statistical Analysis

All statistical analyses were performed with the Statistical Package for the Social Sciences (SPSS) software (IBM SPSS Statistics for Windows, Version 25.0, IBM Corp, Armonk, NY). Repeated measures within-subjects analysis of variance (RM-ANOVA) was used to assess statistically significant differences between dependent variables (RMSE, CBF, CBV, etc.) obtained from different TTDs. For GM and WM regions, averages of these dependent variables were taken for each subject so they could be considered independent.

The assumption of normality was tested with the Shapiro-Wilk test, where *p* > 0.05 indicated normally distributed data. The assumption of sphericity was tested using Mauchly’s test. In cases of non-sphericity, the degrees of freedom were corrected to decrease the Type-I error rate. For pairwise comparisons, a Bonferroni-corrected *p*-value of 0.008 was used to keep the overall Type-I error at 5%.

When the assumption of normality failed, the non-parametric Friedman’s test was performed. Then, to identify pairwise significance, the Wilcoxon signed rank test was used with Bonferroni-corrected significance values.

## 4. Results

### 4.1. Goodness of fit

Figure 3 (a-d) gives typical fits of estimated signal to a measured GM STC with each TTD. All fits capture the rapid initial drop in signal and accurately follow the recirculation bump. Figure 3 (e-h) shows the corresponding TTDs for each fit. The shape of the gamma-variate TTD marginally differs from those of other TTDs, as does its corresponding MTT estimate. Table 1 gives the means and SDs of RMSE of the fits with four different TTDs averaged over all subjects for GM and WM.

**Fig 3.**
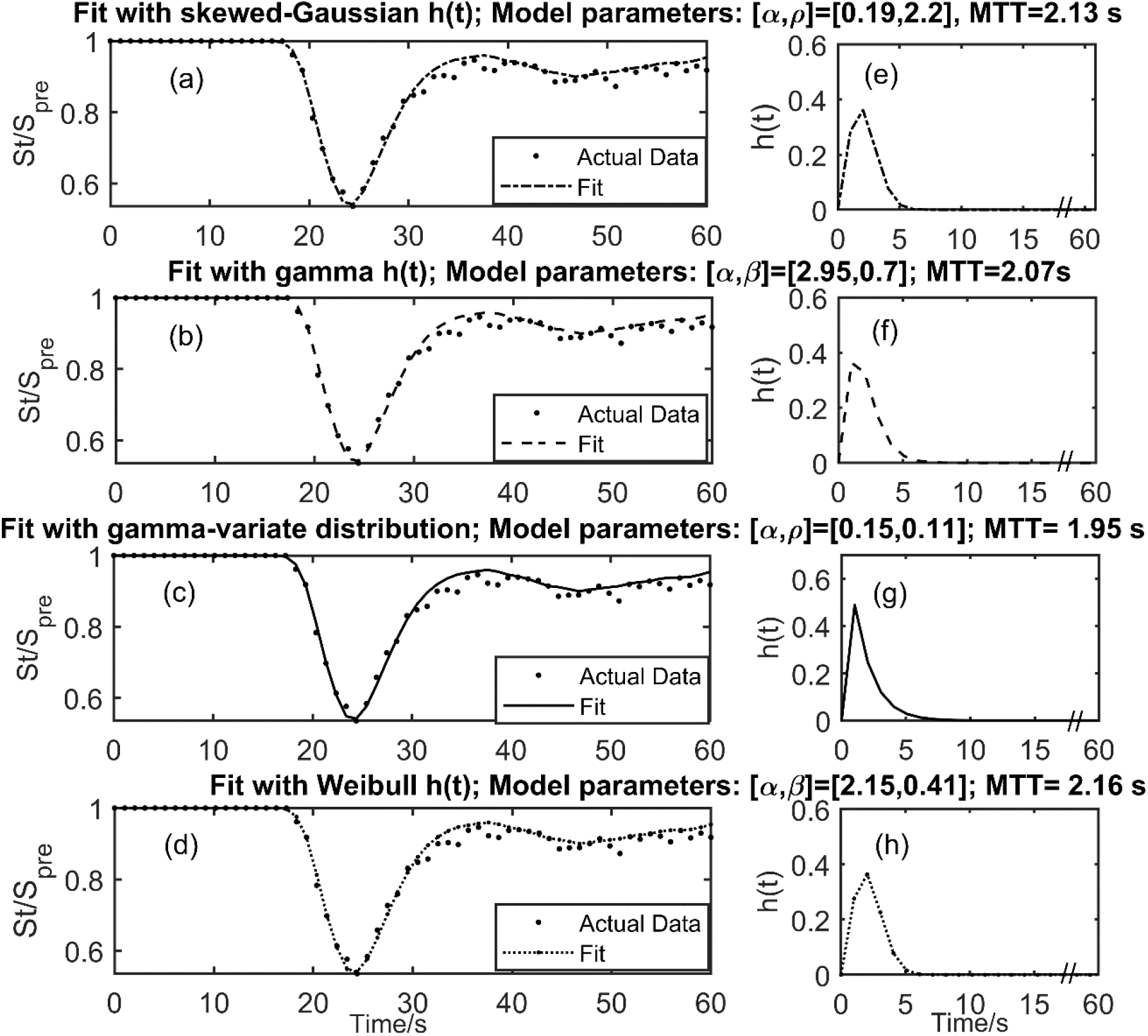
(a-d, left) Dot plots showing typical fits to a baseline-normalized GM signal time curve using skewed-Gaussian, gamma, gamma-variate, and Weibull distributions, respectively. (e-h, right) Corresponding transit time distributions (TTDs) for each fit. Parametric signals fitted well to the measured data. All TTDs, except gamma-variate, have similar shapes and mean transit times (MTTs). Note: The baseline data showed a similar level of noise to post-contrast data; however, for ease of demonstration, all the datapoints before bolus arrival are considered to have the same signal intensity as the mean of the pre-bolus signal intensities, S_pre_

**Table 1:**
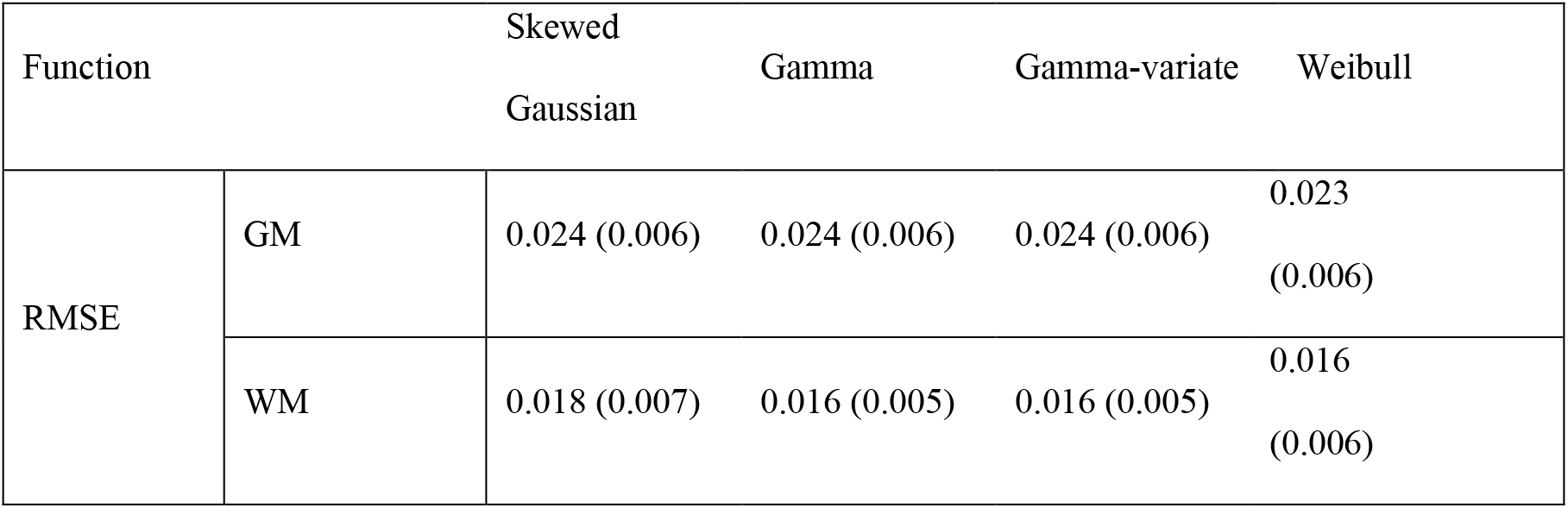
Mean (SD) of root-mean-square error (RMSE) obtained with each transit time distribution (TTD) for grey matter (GM) and white matter (WM)

For the RMSE of GM fitting, the Shapiro-Wilk test indicated normality (*p* > 0.05) and Mauchly’s test indicated non-sphericity (*p* < 0.001). After Greenhouse-Geisser correction of the degrees of freedom, RM-ANOVA revealed no statistically significant differences in the RMSEs obtained using different TTDs (*p* = 0.113).

For the RMSE of WM fitting, the RMSE values deviated from a normal distribution *(p* < 0.001). Friedman’s test indicated no statistically significant differences between the RMSEs (*p* = 0.021).

### 4.2. Parameter Estimates

Table 2 gives estimated mean values and SDs of CBF, MTT, and CBV with each TTD, averaged over all subjects. Perfusion estimates obtained with all four TTDs are comparable to the literature values shown.

**Table 2:**
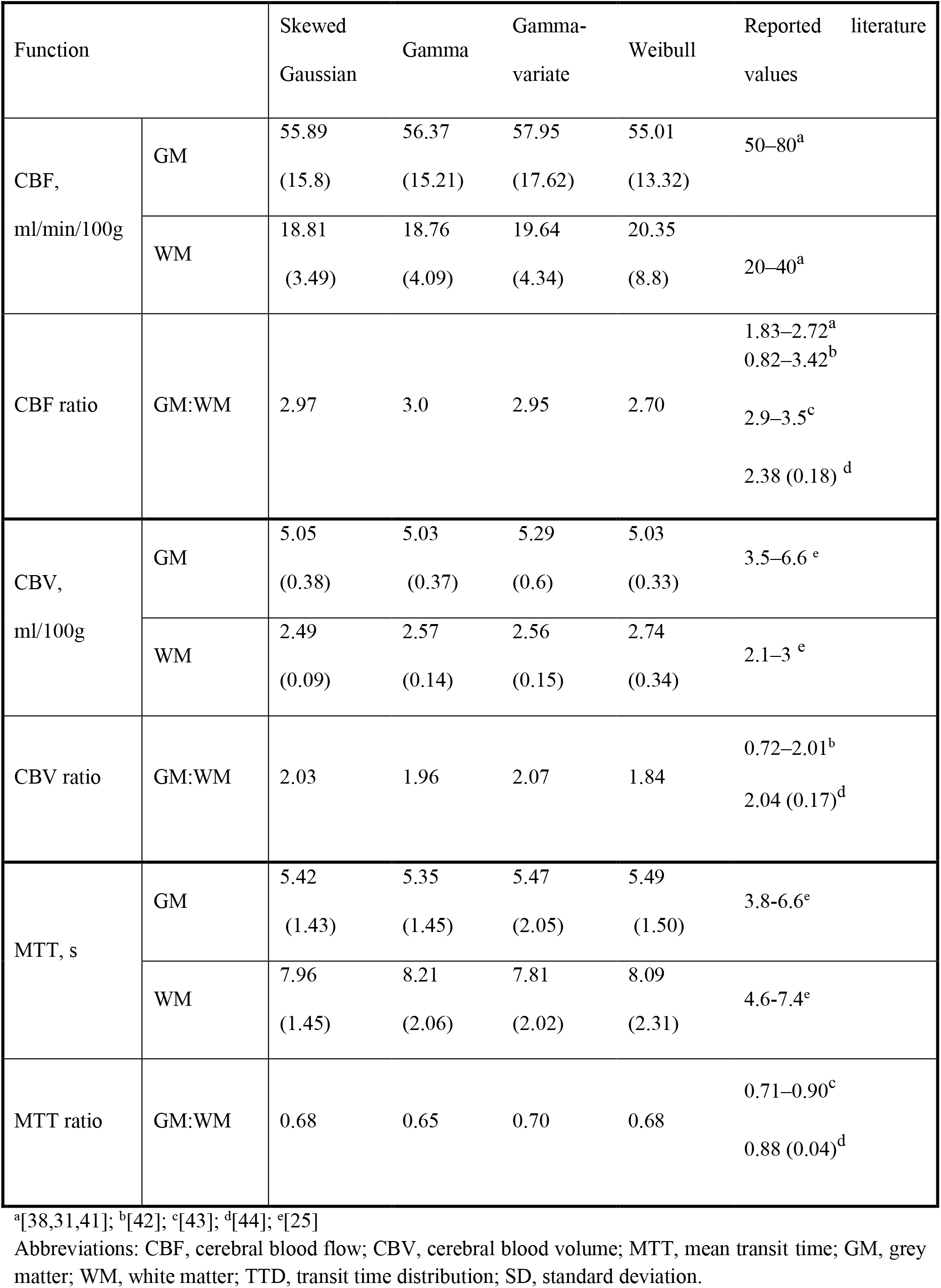
Mean (SD) of parameter estimates obtained with each transit time distribution (TTD), h

For the CBF, MTT, and CBV estimates in GM and WM, the Shapiro-Wilk test indicated normality (*p*_GM_, *p*_WM_ > 0.05) and Mauchly’s test indicated non-sphericity (*p*_GM_ ≤ 0.001, *p*_WM_ ≤ 0.013). After Greenhouse-Geisser correction, RM-ANOVA revealed no significant differences in CBF, MTT, or CBV estimates (*p_GM_* = 0.147, 0.241, 0.851, respectively; and *p*_WM_ = 0.239, 0.052, 0.259, respectively) for GM and WM with four TTDs.

### 4.3. Bias and dispersion

Figure 4 shows the relative bias in CBF, MTT, and CBV estimates for all four models in the GM (top row) and WM (bottom row) for different SNRs. For GM-CBF, the skewed-Gaussian distribution showed the lowest bias over all SNRs. The relative bias in MTT and CBV was the lowest with gamma and the highest with skewed-gaussian TTD. For all models, the bias in GM-MTT and GM-CBV approached an asymptote at an SNR of around 40, with minimal improvement beyond this point. For WM-CBF estimation, gamma was the most accurate and skewed-Gaussian was the least accurate model. For WM-MTT and WM-CBV, gamma-variate and skewed-Gaussian were the least and most accurate models, respectively. Bias in both WM-CBV and WM-MTT was much lower than the equivalent parameters in GM for noisy signals. The relative bias of all models converged at an SNR of 50.

**Fig 4.**
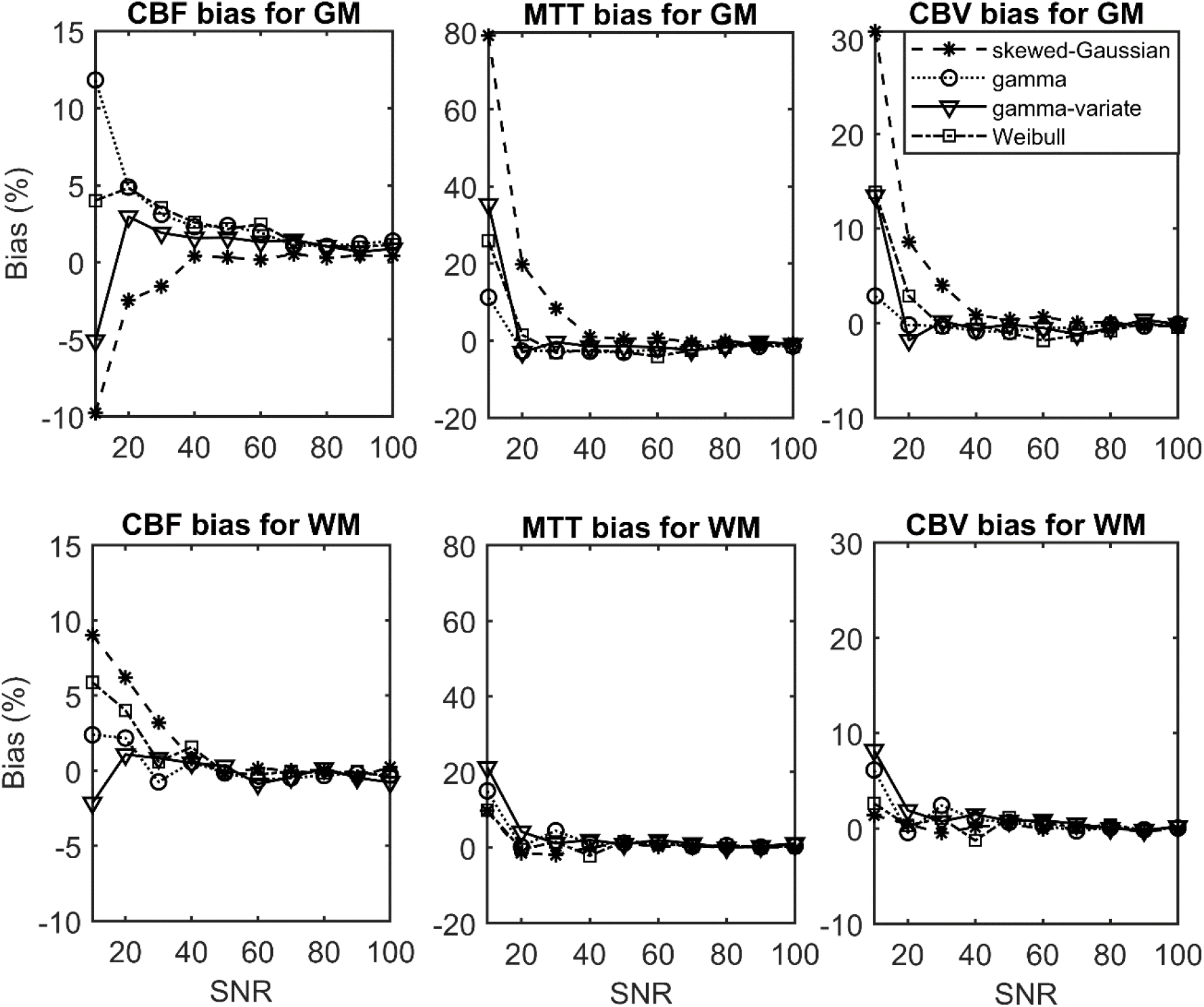
Relative bias in perfusion parameter estimates with four different models of transit time distribution (TTD) as a function of signal-to-noise ratio (SNR). The gamma function gives more accurate parameter estimates than others for noisy gray matter (GM) signals. The accuracy of perfusion estimates in white matter (WM) shows little variation between models, with no improvement after SNR of 40 for either GM or WM. Abbreviations: CBF, cerebral blood flow; CBV, cerebral blood volume; MTT, mean transit time

Figure 5 shows the relative dispersions in CBF, MTT, and CBV estimates for all four models in GM (top row) and WM (bottom row) for different SNRs. All models showed similar trends in their relative dispersions: the spread in the estimates decreased with increasing SNR. Dispersion in GM-CBF was similar for all models; however, for GM-MTT and GM-CBV, the skewed-Gaussian TTD showed more variance, especially in noisy signals. Relative dispersion in GM-CBF, GM-MTT, and GM-CBV was the lowest with the Weibull, gamma-variate, and gamma functions, respectively. For WM, the gamma function gave the most precise CBF and MTT estimates, whereas the skewed-Gaussian function was the best for CBV estimation over all SNRs. The precision of all models tended to converge beyond an SNR of 40.

**Fig 5.**
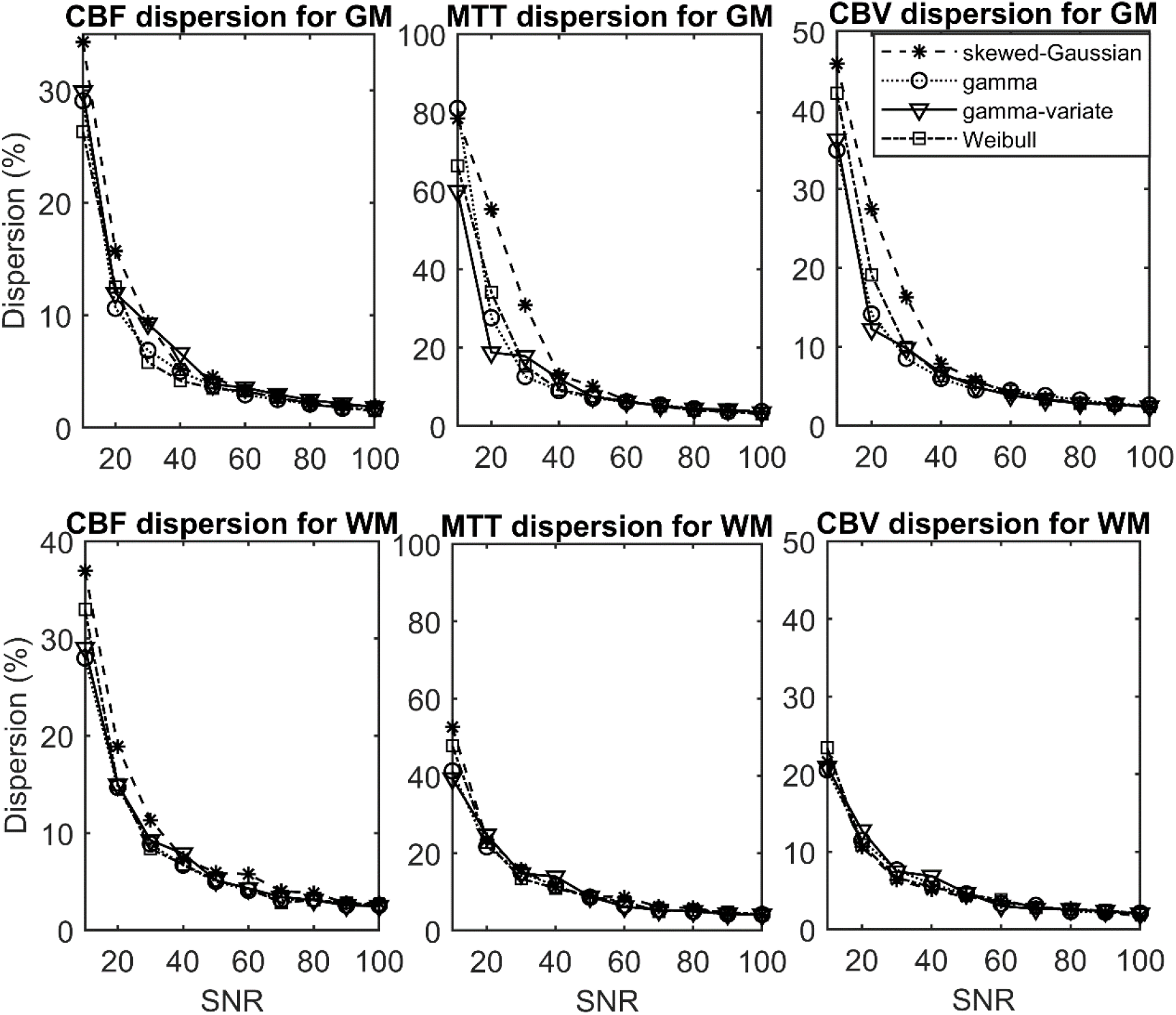
Relative dispersion in perfusion estimates for all four models of transit time distribution (TTD) as a function of signal-to-noise ratio (SNR). All models show similar trends in dispersion: the spread of the parameter estimates decreases with increasing SNR. The gamma and gamma-variate functions tend to perform better for noisy gray matter (GM) signals. Abbreviations: CBF, cerebral blood flow; CBV, cerebral blood volume; MTT, mean transit time.

### 4.4. Success rate

Table 3 presents the means and SDs of the percentage success rate of fitting GM and WM signals with different TTDs. For GM and WM signal fitting, the Shapiro-Wilk test indicated that the success rates were not normally distributed (*p*_GM_, *p*_WM_ < 0.001). Friedman’s test indicated significant differences (*p*_GM_, *p*_WM_ < 0.001) between the success rates for both GM and WM. For GM, fits with the skewed Gaussian and gamma TTDs had significantly higher success rates (*p*_GM_ < 0.008) than those with other distributions. For WM, the gamma distribution had a significantly higher success rate (*p*_WM_ < 0.008) than all other distributions.

**Table 3:**
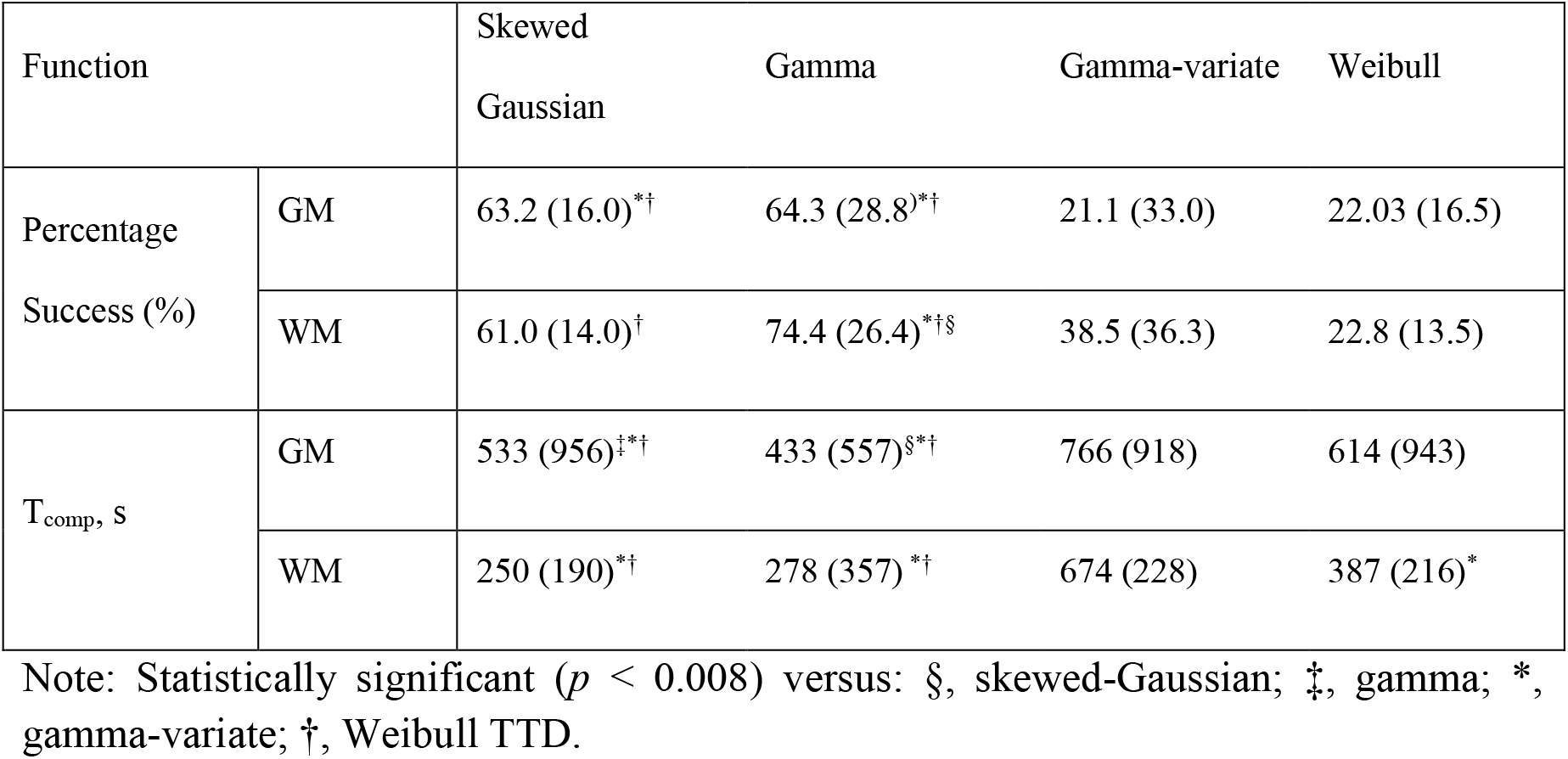
Mean (SD) success rate (percentage of successful fits) and computation time with each TTD.

### 4.5. Computation Time

Table 3 also shows *T_comp_’s* for the four TTDs averaged over all subjects in both GM and WM. It took longer to fit the GM signals than the WM signals, with a larger variation of *T_comp_*. For both GM and WM signal fitting, the Shapiro-Wilk test indicated that the *T_comp_’*s significantly deviated from a normal distribution (*p*_GM_, *p*_WM_ < 0.001). Friedman’s test indicated that there was a significant difference between the *T_comp_’s* (*p*_GM_, *p*_WM_ < 0.001) for different TTDs. For GM, fits using gamma TTDs converged significantly faster (*p_GM_* < 0.008) than all other distributions. For WM, fits with the gamma and skewed Gaussian distributions converged faster than other distributions (*p*_WM_ < 0.008).

## 5. Discussion

The primary aim of this study was to compare the utility of four different forms of TTD for model-dependent deconvolution. We propose the gamma function as the most suitable TTD for perfusion quantification applications, as it provides faster and more stable fits. The seconds saved by the gamma TTD in ROI-based analyses could save many minutes when applied to whole-brain perfusion map generation. This may be particularly useful for clinical cases that require rapid analysis and decision-making, which will in turn facilitate faster diagnosis and initiation of treatment. Moreover, as compared to other TTDs, the perfusion parameters derived from the gamma model are less sensitive to noise and less dependent on the initial guesses. This indicates that the gamma model is likely to generate robust, reproducible perfusion estimates, making it better-suited for cross-center studies.

Curve fits—with all functions—were excellent, with low RMSEs. Perfusion estimates from the four TTDs were also congruent with previously reported values. Since all functions gave similar RMSEs and perfusion estimates, we considered a TTD to be more suitable than another when it provided at least one computational benefit, such as lower sensitivity to noise, higher stability of fit, or lower computation time [9].

Our bias and dispersion analysis demonstrates that the four TTDs studied here differ in terms of the accuracy of their perfusion estimates for low SNRs, but they quickly converged beyond an SNR of around 40-50. In terms of accuracy, out of the six possible combinations (two tissue types, three perfusion parameters), both the gamma and skewed-Gaussian model performed the best in three. So, when only accuracy is considered, either a gamma or skewed-Gaussian TTD can be used, depending on the parameter of interest and tissue type. However, in terms of spread of the estimates (i.e., dispersion), the skewed-Gaussian function gave the least precise estimates in all six possible combinations of parameter and tissue type. The gamma model, on the other hand, gave the most precise estimates in three combinations: GM-CBV, WM-CBF, and WM-MTT. So, if both accuracy and precision are considered, the gamma model performs best. Additionally, our analysis shows that a minimum SNR of 40 is required to produce accurate and precise perfusion estimates in general, with slightly lower SNR limits being possible for certain combinations of parameters and tissues.

This study also indicates that the gamma TTD produces more stable fits; that is, a higher percentage of fits converged to the global minimum. This is likely because the gamma function results in a smoother error surface with fewer local minima than other TTDs. Further, given that it has a higher chance of convergence irrespective of the initial guess, a gamma TTD can be used to accelerate curve-fitting.

In addition to the usual perfusion parameters, the model-based approach taken in this work lends itself to the calculation of further parameters related to the width and shape of the TTDs. Vessels created by tumor angiogenesis are chaotically structured, dilated, and irregularly shaped, so the GBCA particles exhibit a broad range of transit times as they traverse them. Consequently, the TTDs for tumor regions may be wider than those of normal regions and the width and shape of a TTD might be used to distinguish between normal and tumor vessels. Another advantage of the model-dependent deconvolution approach presented here is that it is more robust against experimental noise than model-independent deconvolution [11]. This is because this approach can characterize the residue function using only two free parameters, without estimating it at every time point. Therefore, our ROI-based perfusion measurements can be straightforwardly extended to pixel-wise estimation of perfusion, creating brain maps of CBF, CBV, and MTT. Such maps can provide clinicians with information on capillary flow profiles and allow them to characterize tissue viability in ischemia [6]. Additionally, maps of flow heterogeneity and oxygen extraction fraction can be generated using the MTT and the SD of the TTD respectively [11,23]. Due to its high fit-stability and low time-complexity, the gamma TTD can offer a fast and effective brain map generation for these studies.

There are some limitations to this work. First, the perfusion estimates were not compared directly with estimates obtained by positron emission tomography (PET), the reference standard for perfusion imaging. Second, we did not consider signals from pathological tissues, which are precisely the cases where TTDs are more likely to deviate and thereby produce statistically different perfusion estimates. This is because the primary purpose of the study was to compare different forms of *h,* rather than assess the absolute accuracy of the approach. Future extensions of this work will compare TTDs in pathological tissues where stenosis with marked turbulence, tumors, BBB leakage, or irregular collateral blood-flow paths may account for additional uncertainty [11]. Third, we used a ‘global AIF’ under the premise that it gives a reasonable representation of the arterial input to every ROI. Global AIFs can be delayed and spread (i.e., dispersed) in reaching the ROI. Typically, the arterial dispersion effect is more pronounced in stroke patients. As the present patient cohort had no reported vessel disease, the effect of arterial dispersion was regarded as trivial. Additionally, it has been shown that the flow estimates are independent of vascular delay for model-dependent approaches [11]. Therefore, we did not consider the arterial delay and dispersion of the arterial bolus. Saying that, future extensions of this work can investigate measures of local AIF in the tissue neighborhood [45] or mathematically modelling the effects of delay and dispersion of the AIF [9]. Lastly, the relative bias may have been underestimated here. Some residual noise may still exist in our ground truth—the patient-derived signal with an SNR of 100. While simulations can produce a ground truth that is free of such noise-related bias, we instead elected to use high SNR *in vivo* data, which are more representative of real physiological processes.

## 6. Conclusion

We showed that the gamma distribution is superior to other plausible TTD functions. Although all four functions gave perfusion estimates similar to published studies, the gamma TTD gave estimates that are more robust against noise. Moreover, the gamma TTD offers significantly faster convergence with higher stability of fit. Therefore, it can substantially decrease the overall computation time and perform reliably under noisy conditions.

## Abbreviations

AIF: Arterial input function
ANOVA: Analysis of variance
AU: Arbitrary units
AUC: Area under the curve
AV: Arterial voxel
CBF: Cerebral blood flow
CBV: Cerebral blood volume
CSF: Cerebrospinal fluid
CTC: Concentration time curve
DCE-MRI: Dynamic contrast-enhanced-MRI
DSC-MRI: Dynamic susceptibility-contrast-MRI
FM: First moment
FWHM: Full width at half maximum
GBCA: Gadolinium-based contrast agent
GRE: Gradient recalled echo
MTT: Mean transit time
RI: Roughness index
RMSE: Rootmean-square error
ROI: Region of interest
SD: Standard deviation
SNR: Signal to noise ratio
SPSS: Statistical package for the social sciences
STC: Signal time curve/course
SVD: Singular value decomposition
TTD: Transit time distribution.

## Supporting Information

Simulation of different forms of transit time distributions Figure 1 shows the shape of four TTDs in three different vascular conditions. We simulated three STCs mimicking three different tissues: healthy GM, ischemic (internal carotid stenosis) [22], and tumor (high grade glioblastoma multiforme) [21] as follows. Gamma function was used as the underlying TTD [6,21,22]. The shape and scale parameters for these three conditions were taken from published studies of Schabel [21], Larsson et al. [22], and Vonken et al. [32](given in Supporting Table 1). With these gamma TTDs, three unique *R*s were derived using equation (2). The CTCs were created using equation (1) with AIF obtained by aligning and averaging the participant AIFs and CBF values representative of the three vascular conditions [21,22,31,32] (shown in Supporting Table 1). Finally, CTCs were converted to STCs by using a non-linear relationship [27].

To visualize how the skewed gaussian, gamma-variate, and Weibull TTDs characterize different vascular conditions, the parametric signals derived from them were fitted to the three tissue signals created above. The results are shown in Figure 1.

**Supporting Table 1:**
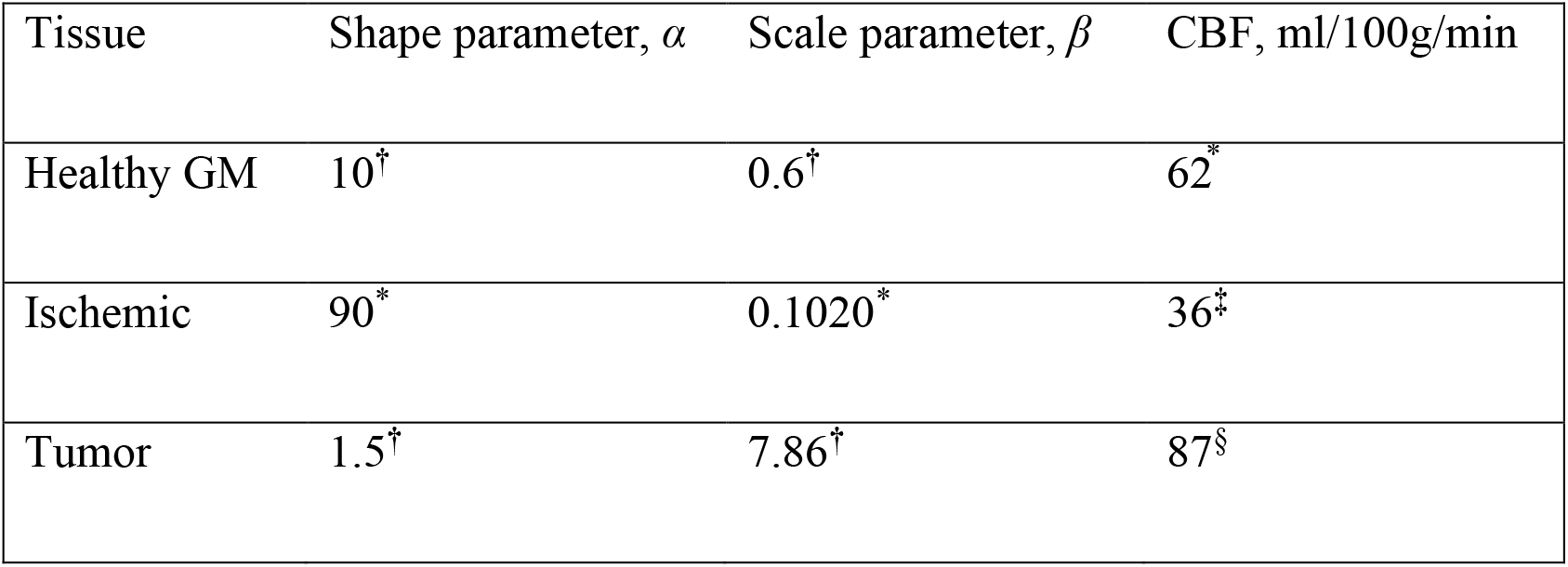
Shape and scale parameters of gamma TTD with cerebral blood flow (CBF) values for simulating healthy and pathological signals, taken from studies by: †) Schabel [21]; *) - Larsson et al. [22]; and §) Vonken et al. [32].

## 7. Author’s Contribution

Sobhan: Analysis and interpretation of data, drafting of manuscript. Gkogkou: Critical revision Johnson: Study conception and design, acquisition of data, critical revision. Cameron: Study conception and design, interpretation of data, critical revision.

